# Deep trans-omic network fusion reveals altered synaptic network in Alzheimer’s Disease

**DOI:** 10.1101/2022.05.02.490336

**Authors:** Linhui Xie, Yash Raj, Pradeep Varathan, Bing He, Kwangsik Nho, Shannon L. Risacher, Paul Salama, Andrew J. Saykin, Jingwen Yan

## Abstract

Multi-omic data spanning from genotype, gene expression to protein expression have been increasingly explored to interpret findings from genome wide association studies of Alzheimer’s disease (AD) and to gain more insight of the disease mechanism. However, each -omics data type is usually examined individually and the functional interactions between genetic variations, genes and proteins are only used after discovery to interpret the findings, but not beforehand. In this case, multi-omic findings are likely not functionally related and therefore give rise to challenges in interpretation. To address this problem, we propose a new interpretable deep neural network model MoFNet to jointly model the prior knowledge of functional interactions and multi-omic data set. It aims to identify a subnetwork of functional interactions predictive of AD evidenced by multi-omic measures. Particularly, prior functional interaction network was embedded into the architecture of MoFNet in a way that it resembles the information flow from DNA to gene and protein. The proposed model MoFNet significantly outperformed all other state-of-art classifiers when evaluated using multi-omic data from the ROS/MAP cohort. Instead of individual markers, MoFNet yielded multi-omic sub-networks related to innate immune system, clearance of misfolded proteins, and neurotransmitter release respectively. Around 50% of these findings were replicated in another independent cohort. Our identified gene/proteins are highly related to synaptic vesicle function. Altered regulation or expression of these genes/proteins could cause disruption in neuron-neuron or neuron-glia cross talk and further lead to neuronal and synapse loss in AD. Further investigation of these identified genes/proteins could possibly help decipher the mechanisms underlying synaptic dysfunction in AD, and ultimately inform therapeutic strategies to modify AD progression in the early stage.

## Introduction

Alzheimer’s Disease (AD) is a complex, heterogeneous brain disorder, with high heritability indicating substantial genetic risk. Large scale genome wide association studies (GWASs) have revealed a significant number of genetic variants in association with AD and related traits, e.g., single nucleotide polymorphism (SNPs) in *APOE, BIN1, CLU, CR1* and *PICALM* [1–7]. However, the downstream biology through which these SNPs contribute to the risk of disease development is still not fully understood [8–10]. It remains a major challenge to convert GWAS findings to targetable mechanism for further drug development and therapeutic intervention.

To further understand these GWAS findings, multi-omic data have been increasingly explored spanning from genotype, gene expression to protein expression [11, 12]. Integrative - omics analysis is expected to identify the genetic markers evidenced from multiple aspects and provide more insights of molecular mechanisms underlying AD. Early multi-omic studies examined multi-omic data in an isolated fashion or in a sequential way. For the first type, each -omics data type is analyzed individually and then overlapped genetic markers across data types are extracted as hits of top interest [13]. For the second type, findings from upstream layers (e.g., genotype) were used as seed to narrow down the search space in the downstream layers (e.g., gene expression) [14, 15]. In either scenario, functional connections between -omics data types are minimally considered. They have very limited power to detect genetic markers that do not directly overlap across -omics data types but instead are functionally related [16–18].

To address these challenges, we developed an interpretable deep learning method, multi-omic fused neural network (MoFNet), to jointly model multi-omics data and the prior functional interactions between genetic variations, genes and proteins (e.g., eQTLs and transcriptor binding effect). To achieve interpretability, prior functional interaction across data types was embedded into the neural network architecture to mimic the information flow from DNA to gene and protein. That said, we used prior functional interactions to define the biological architecture of MoFNet, compared to the conventional fully connected ‘black box’ neural networks. MoFNet can also simultaneously prioritize variants and molecular mechanisms such as gene regulation and lead to a sub-network of SNPs, genes and proteins predictive of AD outcomes. MoFNet was applied to the genotype, gene expression and protein expression data from the ROS/MAP cohort, aiming to return a multi-omic subnetwork for better understanding of disease mechanism and to identify possible intermediate molecular targets for therapeutic intervention.

## Material and Methods

### Religious Orders Study and Memory and Aging Project (ROS/MAP)

Multi-omic data used for discovery were obtained from the Religious Orders Study (ROS) and Memory and Aging Project (MAP) cohorts [19]. Both cohorts aim to study the risk factors for cognitive decline and incident AD dementia, and have been providing valuable multi-omic data resource to the research community. We downloaded imputed genotype, RNA-Seq (ribonucleic acid sequencing) gene expression, protein expression and diagnosis information for ^-^600 participants via Accelerating Medicines Partnership for Alzheimer’s Disease (AMP-AD) portal [20]. All the gene expression and protein expression were collected from prefrontal cortex region from postmortem brains of participating subjects. Subjects were excluded from the subsequent analysis if 1) they miss one or more -omics data types, or 2) are part of AD dementia group but with other cause of cognitive impairment. Finally, 133 subjects with full set of three -omics data types were included (Table 1). Detailed preprocessing steps of genotype, RNA-Seq and protein expression data can be found in the supplementary sections S.1, S.2, and S.3.

**Table 1.** Participant demographic information.

| Diagnosis | ROS/MAP |  | MSBB |  |
| --- | --- | --- | --- | --- |
|  | CN | AD | CN | AD |
| Subject Number | 77 | 56 | 41 | 80 |
| Male/Female | 35/42 | 22/34 | 21/20 | 23/57 |
| Education(mean± std.) | 16.7 ± 3.2 | 16.8 ± 3.7 | N/A | N/A |
| Age(mean± std.) | 83.0 ± 4.5 | 86.3 ± 3.5 | 83.3 ± 8.1 | 85.1 ± 6.8 |

### Mount Sinai Brain Bank (MSBB)

The replication data set was from Brodmann 10 region of the Mount Sinai Brain Bank (MSBB) [21]. This cohort was composed after strict inclusion/exclusion criteria, representing the full range of cognitive and neuropathological disease severity, without significant non-AD neuropathology. All participants underwent neuropathological evaluation according to the Consortium to Establish a Registry for Alzheimer’s Disease (CERAD) protocol [22]. In this study, we included 121 white participants with genotype, RNA-Seq and protein expression data. RNA-Seq data of MSBB and ROS/MAP participants both went through the same pre-processing pipeline, adjusted for age, gender, education, batch and RNA integrity number (RIN) as covariates, and were directly downloaded as quality controlled from the AMP-AD knowledge portal. Detailed preprocessing steps of genotype, RNA-Seq and protein expression data can be found in the supplementary sections S.1, S.2, and S.3.

Shown in Table 1 is the detailed demographic information of all participants included from the ROS/MAP and MSBB cohorts. Female/male ratio is relatively higher in AD group than in cognitive normal (CN) group and AD group is on average 2-3 years older in both cohorts. This is consistent with existing findings that gender and age are two prominent risk factors for AD [23].

### SNP and Gene Filtering

Given the large number of SNPs and genes, it is infeasible to directly model all genetic variants and evaluate their overall predictive power. Considering that protein expression is the data bottleneck in ROS/MAP with only 186 peptides (mapped to 126 unique genes), we took a bottom-up approach to narrow down the total number of -omics features by using these peptides as seeds and selecting only a subset of SNPs, genes functionally related to them (Fig. 1). In the proteomic layer, abundance level of 186 peptides were measured in the ROS/MAP cohort. These peptides were mapped to 126 unique genes (gene set A), which were found to interact with 954 genes (gene set B) in the functional interaction network obtained from the Reactome database [24]. Among these 1080 (126 + 954) genes, 743 without missing RNA-seq data were included to represent the transcriptomic layer in our model. In the genomic layer, we identified SNPs located on the upstream of these 743 genes within the boundary of 5K. To ensure the functional connection of selected SNPs and their downstream genes, we included only SNPs significantly associated with the transcription factor-binding activity, based on the single nucleotide polymorphisms to transcription factor binding sites (SNP2TFBS) database [25]. Taken together, we have 822 SNPs, 743 genes and 186 peptides for the subsequent modeling. The functional relationships used to filter the genes and SNPs form a trans-omic functional interaction network, which will be embedded into the architecture of deep neural network to guide the search of functionally connected features related to AD (Fig. 1 (c)).

**Fig. 1.**
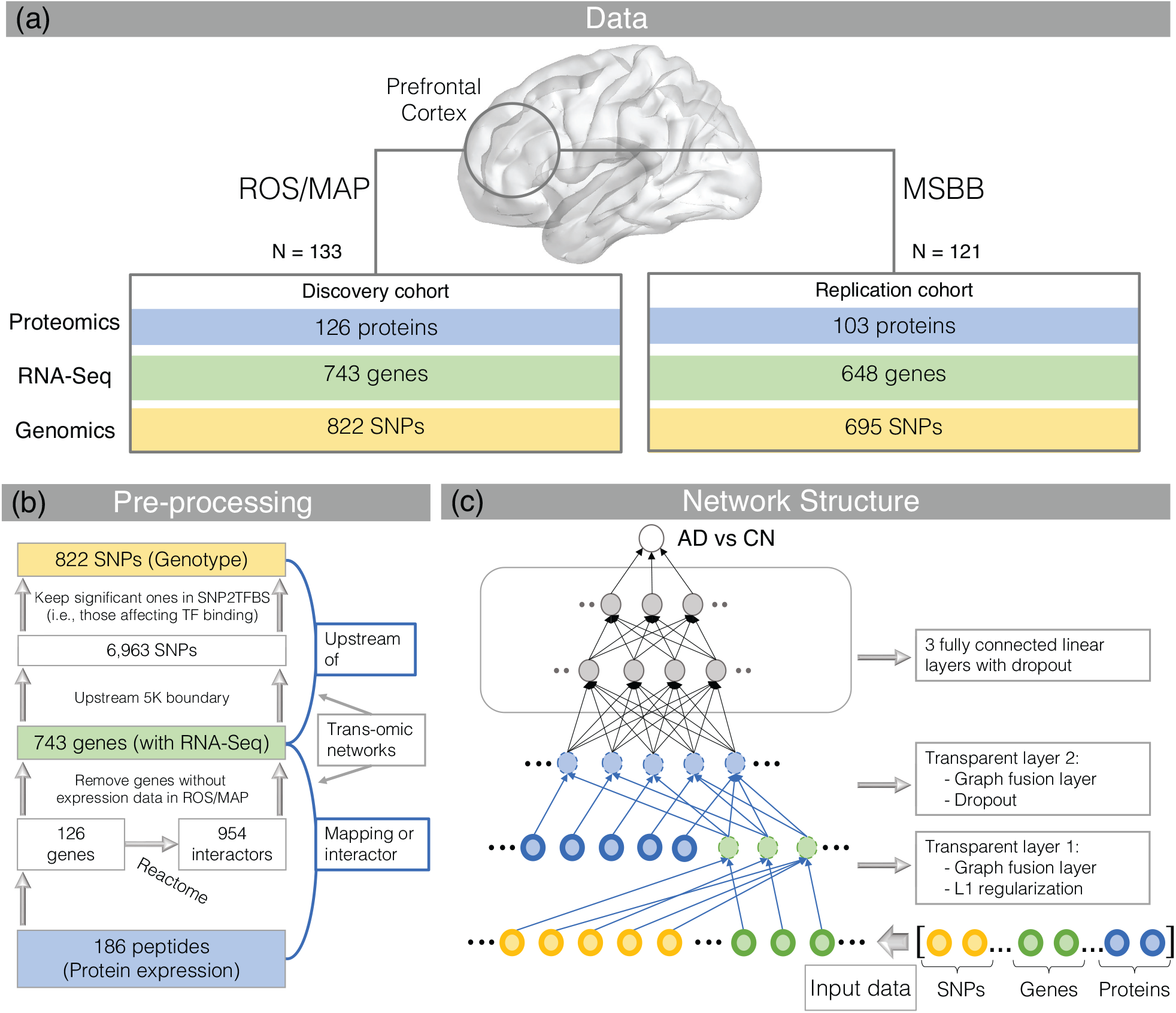
Multi-omic data used, filtering strategy and the architecture of MoFNet. (a) Multi-omic data in discovery and replication cohorts. (b) SNP and gene filtering in the ROS/MAP cohort using peptide as seed. (c) Architecture of the proposed MoFNet model, with the prior trans-omic network in (b) embedded in the first two transparent layers.

### Prediction Outcome

Extracted SNP genotype, gene expression and protein expression data were used to classify AD patients from cognitive normals (CNs). For all the participants included in this study, their final clinical diagnosis at the point of brain tissue collection was used to indicate their disease status. In this case, the diagnosis time remains consistent with the expression data collection time.

### Architecture of MoFNet

The proposed deep multi-omic fused neural network model (MoFNet), extended from Varmole [26], is a graph neural network with two transparent layers structured based on the functional connectivity between selected SNPs, genes and proteins (i.e., trans-omic network shown in Fig. 1(b)). Shown in Fig. 1(c) is the detailed architecture of MoFNet. The first transparent layer has 1565 input nodes, corresponding to 822 SNPs and 743 genes respectively, and 743 output nodes, corresponding to 743 enhanced genes. Links in the first transparent layer were added if one SNP (as an input node) is connected to one gene (as an output node) in the prior transomic network. In addition, we added links between duplicated genes nodes (i.e., gene A node in the input and gene A node in the output). In this case, output gene nodes will have enhanced expression data by integrating information from its upstream SNPs and its originally measured expression level. We assume that not all SNPs are equally informative and helpful in enhancing the signal of the downstream gene. Therefore, L1 regularization was applied for the first transparent layer such that links between non-informative SNPs and their downstream genes will mostly get zero weight. For the second transparent layer, we have 929 input nodes, corresponding to 743 enhanced genes and 186 peptides, and 186 output nodes, corresponding to 186 enhanced peptides. Links were added if one gene (as an input node) is connected to one peptide (as an output node) as indicated in the prior trans-omic network. We also added links between the duplicated peptide (i.e., peptide A node in the input and peptide A node in the output). Taken together, the enhanced peptide nodes, as the output of the second transparent layer, will integrate the information from its corresponding genes, their interactors and upstream SNPs. After that, we have 3 fully connected layers to classify the AD patients from cognitive normal subjects. We used dropout and early termination to avoid overfitting.

1. Input *X*_1_ is the concatenation of the gene expression matrix *G*^*n*×*g*^ (*n* samples by *g* genes), and SNP genotype matrix *S*^*n*×*s*^ (*n* samples by *s* SNPs). 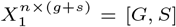 where [·] stands for row concatenation.
2. The output from the first transparent layer *Z*_11_ has the dimension as the number of genes *g*. Links in this layer indicate the prior functional connections between SNPs and genes, and between same genes. Functional connections between SNPs and genes were encoded in an adjacency matrix 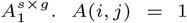 indicates SNP *i* is located upstream of gene *j* and likely to affect the transcription factor binding activity; *A*(*i,j*) = 0 otherwise. We also added self-connections to genes by adding another adjacency matrix 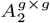, which is an identity matrix with *A*_2_(*i,i*) = 1. Taken these two adjacency matrices together, the first transparent layer is a ‘Biological DropConnect’ layer [26, 27]. Therefore, weight matrix of this layer *W*_1_ has a sparse structure with a dimension of (*s* + *g*) × *g*. Output of this layer 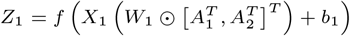 where ⊙ is the Hadamard product, and (·)^*T*^ is the matrix transpose operator.
3. The second transparent layer resembles the structure of the second part of the prior trans-omic network, i.e., the functional connections between genes and proteins. The input of this layer is the concatenation of the protein expression (e.g. peptides) data 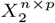 (*n* samples by *p* peptides) and output of the first transparent layer *Z*_11_, i.e., *Z*_1_ = [*X*_2_, *Z*_11_]. The output of the second transparent layer *Z*_2_ has a dimension of the number of peptides. Weight of this layer *W*_2_ has a dimension of (*g* + *p*) × *p*. The adjacency matrix 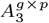 indicates the functional connections between genes and proteins, where *A*_3_(*i,j*) = 1 if gene *i* encodes protein *j* itself or the functional interactor of protein *j*; *A*_3_(*i,j*) = 0, otherwise. Similarly, we added self-connections between peptides with an identity adjacency matrix 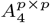 where *A*_4_(*i,i*) = 1. The output of the second transparent layer is 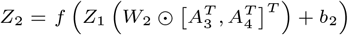.
4. Three fully connected hidden layers *Z*_l_ index by *l* ∈ {3 …*L* – 1} were used together with a sigmoid function in the last layer. *Z_L_* = *σ* (*Z*_*L*–1_ *W_L_* + *b_L_*).
5. Finally, we use binary cross-entropy loss to quantify the classification error: 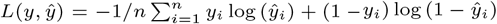.

Due to the small sample size, 5-fold cross validation was applied with grid search to tune the parameters. Details of parameter tuning and final parameters can be found in supplementary section S.4.

### Interpretability

MoFNet can be interpreted in two ways. First, with L1 regularization, weight learned for the links in the first two transparent layers has many zeros. Mapping those weights to the prior trans-omic network will help prune the prior network and lead to sub-networks indicating important information flow from DNA to protein that contributes to the prediction of AD. Secondly, integrated gradient was applied to prioritize node importance for the outcome [28, 29]. The gradient of model prediction for each multi-omic feature indicates how the prediction outcome responds to the changes of the multi-omic features. This importance score will provide the potential explanations in terms of which part of the pruned trans-omic sub-network contribute the most to the disease outcome.

## Results

### Classification Performance

We compared the performance of MoFNet with random forest and four other logistic regression based classification models, using modularity, elastic net, GraphNet and Lasso as penalty terms respectively [30–33]. These sparse logistic regression models were selected because they are designed for both classification and feature selection. Classic classification models, such as support vector machine (SVM) and k nearest neighbor (KNN), do not select features, and therefore are not included for comparison. For the proposed MoFNet, we also evaluated its isoforms structured using shuffled prior trans-omic networks. In total, we repeated shuffling three times and reported the average performance. Two different shuffling strategies were applied: without and without connections between duplicated gene and protein nodes. Shuffling without self connections yielded consistent bad performance and thus are not reported here. The modularity constrained logistic regression was implemented in MATLAB [34]. GraphNet was implemented using R package [35]. Elastic net constrained logistic regression, traditional logistic regression with lasso penalty, and random forest were implemented using the Python scikit-learn package [36]. To provide an unbiased comparison, partition of subjects in all training and testing set was kept identical for all methods. Grid search and 5-fold cross validation for all methods were used to select optimal parameters.

Shown in Table. 2 is the performance of MoFNet and other competing methods on test data set. Due to imbalanced case and control numbers, we reported not only accuracy and AUC, but also F1 score, precision, and specificity metrics to give a comprehensive comparison of performance. In particular, F1 score combines precision and sensitivity(recall) into a single metric, and has been widely used as a major evaluation metric for imbalanced data sets. The proposed MoFNet consistently outperforms other competing models, with highest accuracy, specificity, AUC, and F1-score, indicating its capability in handling the imbalanced data set. The MoFNet structured with shuffled trans-omic network scored the second-best performance. This suggests that the original measures of genotype, gene and protein expression are still the major contributors. The information from functionally connected neighbors indeed provided extra information that may be missed when taking the snapshot of gene and protein expression.

**Table 2.**
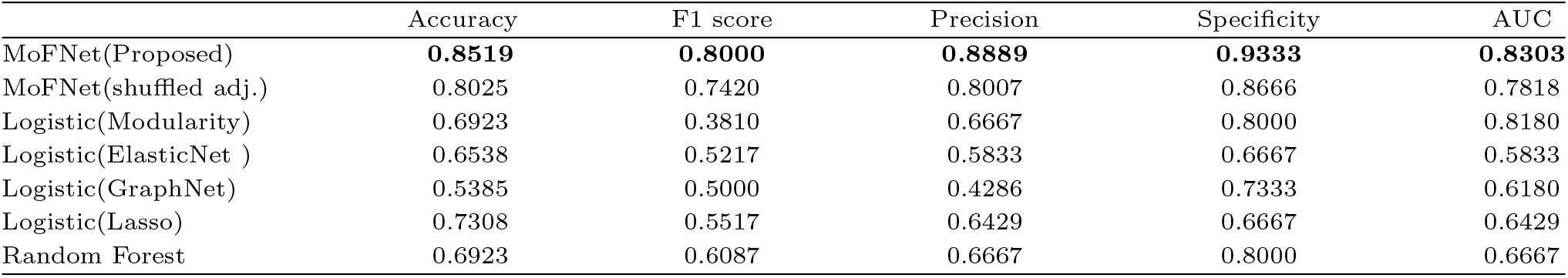
Performance comparison of MoFNet and other competing classification methods across five test data sets (mean ± std).

|  | Accuracy | F1 score | Precision | Specificity | AUC |
| --- | --- | --- | --- | --- | --- |
| MoFNet(Proposed) | **0.8519** | **0.8000** | **0.8889** | **0.9333** | **0.8303** |
| MoFNet(shuffled adj.) | 0.8025 | 0.7420 | 0.8007 | 0.8666 | 0.7818 |
| Logistic(Modularity) | 0.6923 | 0.3810 | 0.6667 | 0.8000 | 0.8180 |
| Logistic(ElasticNet ) | 0.6538 | 0.5217 | 0.5833 | 0.6667 | 0.5833 |
| Logistic(GraphNet) | 0.5385 | 0.5000 | 0.4286 | 0.7333 | 0.6180 |
| Logistic(Lasso) | 0.7308 | 0.5517 | 0.6429 | 0.6667 | 0.6429 |
| Random Forest | 0.6923 | 0.6087 | 0.6667 | 0.8000 | 0.6667 |

### Multiomic sub-networks for AD

In addition, we examined which SNPs, genes and/or proteins contribute to the final prediction and how they are functionally connected. For all competing methods, only modularity and elastic net constrained logistic regression models yielded a subset of SNPs, genes and proteins with a few known functional connections in the prior network. Multi-omic features identified by other competing methods mostly scattered around the prior network with little known connections.

MoFNet returned the importance score for each SNP, gene and protein node, and the weight for each link in the prior trans-omic network. They are used to prune the prior network and generate sub-networks relevant to AD. A set of ordered threshold was examined and the one that lead to the convergence in the number of connected components was selected as the final cut-off value (Supplementary Figure S1). That says, additional features included by going beyond the selected threshold largely are not functionally connected to those top ones. Final pruned trans-omic network has 169 multi-omic features, including 32 proteins, 80 genes and 57 SNPs. These features formed 3 major connected components (i.e., subnetworks) Fig. 2. Node size is proportional to its importance score. The larger the node size, the more contribution it makes to the final prediction. Similarly, edge width is proportional to the weight taken from transparent layers. The thicker the edge is, the more information integration occurs from SNPs to genes or from genes to proteins. As expected, nodes with top importance scores are mostly proteins since they integrated information from SNPs and genes. One peptide of Apolipoprotein E (*APOE*), a strong risk factor for AD [37], is also identified but are not part of top 3 connected componnents. It is likely due to its limited functional connections in our prior network.

**Fig. 2.**
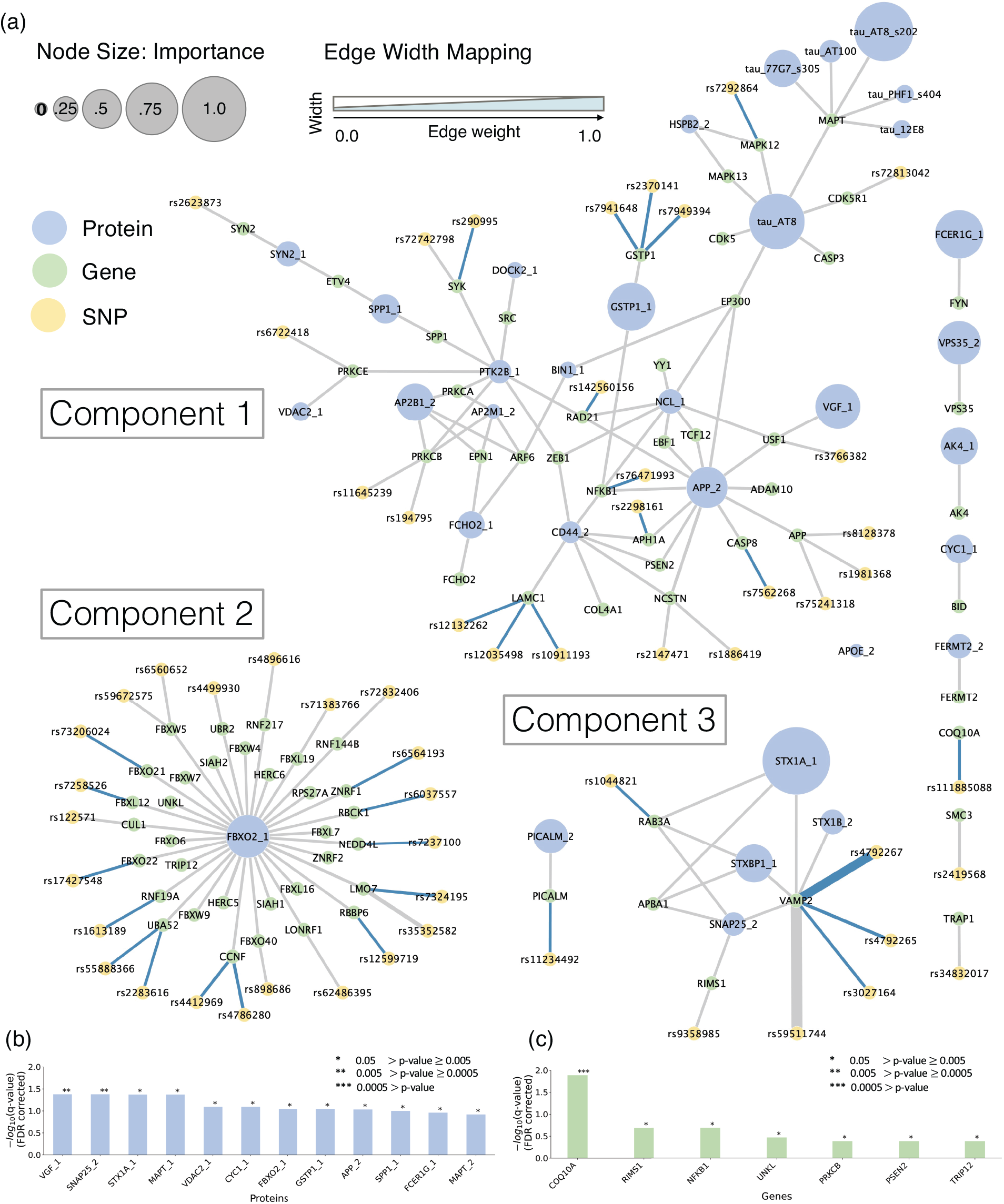
(a) Sub-networks (# of nodes ≥ 2) from pruned trans-omic network using the edge weight and node importance score learned from MoFNet. Node size is proportional to importance score. Edge width is proportional to the edge weight. Blue edges: SNP-gene pairs where the SNP is a known eQTL of its connected gene in the frontal cortex tissue. Numbers at the end of protein indicate different peptides. (b) 12 peptides in (a) with significant differential expression between AD and CN after FDR correction. (c) 7 genes in (a) with with significant differential expression between AD and CN after FDR correction.

The largest connected component has 24 SNPs, 34 genes and 21 proteins, including the amyloid precursor protein *APP* and its corresponding gene. In particular, peptide *APP_2* has the largest connected degree, and peptide *Tau_AT*8 has a large importance score indicating its major contribution to the predicted outcome of each subject. AT8 and AT100, shown at the end of Tau peptides, are antibodies that recognize Tau protein phosphorylated at Ser202/Thr205 and Thr212/Ser214 respectively [38, 39]. Our result also identified other phosphorylated forms of Tau to be highly predictive of AD, such as those recognized by antibodies PHF1, 12E8 and 77G7 [40, 41]. Several studies have demonstrated that the presence of phosphorylated Tau detected by AT8 or AT100, is strongly correlated with the presence of neurofibrillary tangles in AD brains [42, 43]. The activation of the *NLRP3* inflammasome is suggested to contribute to the development of Alzheimer’s disease by promoting the formation of neurofibrillary tangles and the accumulation of abnormal *tau_AT*8 in the brain [44].

### Differential Expression Analysis

For all genes and proteins in the pruned trans-omic network, we further performed differential expression analysis. Since both gene expression and peptide data were pre-adjusted for covariates already, simple unpaired t-test was applied on each gene and peptide respectively. Across 33 peptides and 80 genes identified by MoFNet, 12 peptides and 7 genes showed significant differential expression between AD and CN with FDR corrected *p* ≤ 0.05 (Fig. 2(b) and (c)). Out of those, 7 peptides and 3 genes are from the largest component. Top significant peptides and genes, such as *VGF_* 1, *SNAP25_2* and *COQ10A*, were identified by MoFNet, but not in those major connected components due to their limited functional connections in the prior network. MoFNet returned a lot more genes and proteins since they are either significantly associated with AD themselves or functionally connected to those significant ones (e.g., SNPs or genes), which gives rise to the AD-related sub-networks instead of individual markers.

### Expression quantitative trait locus (eQTL) Analysis

We investigated the function of all SNPs in 3 major components on the downstream transcriptome level. In the Brain eQTL Almanac (BRAINEAC) database, 50 out of 53 SNPs were found significant as eQTL in the frontal cortex region. The rest 3 SNPs were not found. That means, variations in these SNPs are associated with gene expression levels. Further, we examined whether those SNPs are eQTLs of their directly connected genes in Fig. 2. Among all 50 SNP-gene pairs in Fig. 2(a), 29 of them have been identified to have significant associations in the frontal cortex tissue (edges highlighted in blue in Fig. 2). Top significant associations between SNPs and genes from largest component were listed in Supplementary tables: Table S1.

### Enriched cell types

Despite limited single cell multi-omic data in AD, we can still estimate the cell types related to altered genes and proteins from bulk data since each cell type typically has its own marker genes. Those marker genes usually have selectively high expression in specific cell types, but not in others. Based on CellMarker database [45], we examined all the genes and proteins from 3 major network components to check whether they are marker genes of any cell type. Shown in Supplementary Figure S2, genes/proteins identified by MoFNet are mostly marker genes for astrocyte, microglia and neuronal cells. Detailed list of marker genes in each component is shown in Supplementary tables: Table S2. Astrocytes are known to provide support and nourishment to neurons, while microglia cells protect against threats and remove damaged cells [46]. Both astrocytes and microglia are glial cells mediating the neuroinflammation, which has been viewed as a ”double-sword” in AD [47]. Cross-talk between astrocyte and microglia has been recently suggested as a potential target for therapeutic intervention in AD [48].

### Pathway Enrichment Analysis

For all the genes and proteins in 3 major network components, we performed enrichment analysis in REACTOME pathways using g:Profiler [24, 49]. Shown in Fig. 3 is the map of all enriched REACTOME pathways, falling into eight functional groups [50, 51]. Top pathways enriched by genes and proteins in network component 1 include notch1 nlr (NOD-like receptors) signaling, eph ephrin cells, release neurotransmitter cycle, and caspase mediated cleavage. NLR signaling pathways are known to be associated with the inflammatory response in AD [52]. Emerging evidence suggests that Eph-Ephrin signaling is associated with both synaptic dysfunction and immune dysregulation, which in turn promotes the progression of AD [53]. Dysregulation of neurotransmitter release has been involved in the development of AD by affecting neural communication and plasticity, which will possibly lead to the death of nerve cells [54]. Lastly, caspase-mediated cleavage of Tau is viewed as an early pathological event triggering tangle pathology as a critical toxic moiety underlying neurodegeneration [55, 56].

**Fig. 3.**
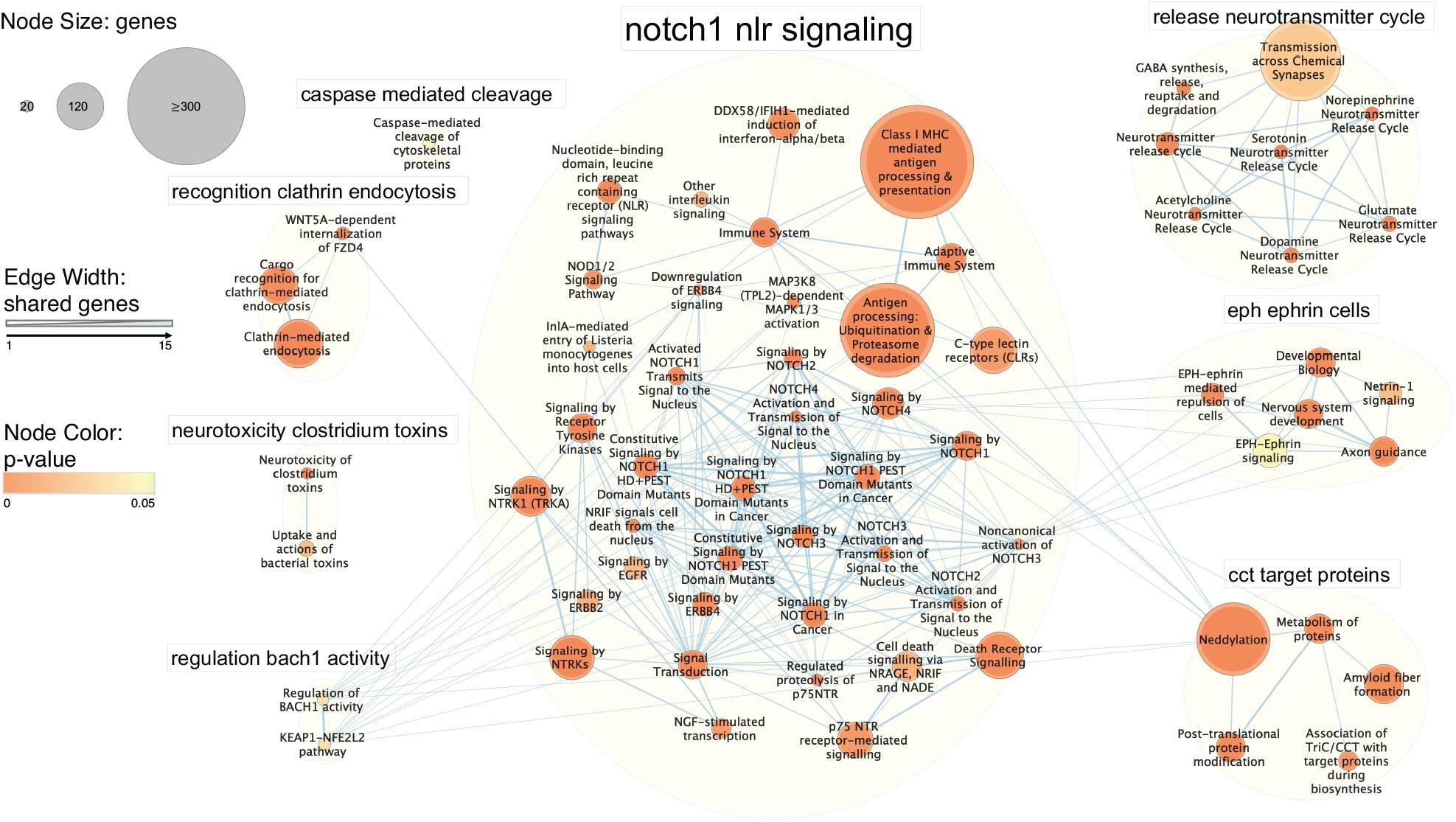
Map of REACTOME pathways enriched by all the genes and proteins identified by MoFNet, toally forming 8 functional groups. Each node is a pathway and node size is proportional to the number of member genes. Edge width is proportional to the number of genes shared between pathways. Node color indicates the significance of each pathway.

The second largest connected component is a sub-network centered around protein *FBXO_2.* It is found closely related to ubiquitination & proteasome degradation as part of antigen processing (adjusted p=1.808e-51) [24, 49]. Neuronal death in Alzheimer’s diseases has a strong connection with misfolded proteins that aggregate within the brain, e.g. Amyloid and Tau tangles. Ubiquitination & proteasome degradation is one of the two major pathways that help get rid of unwanted cells or misfolded proteins to prevent their accumulation and to maintain the health of a cell [57].

The third component is a small sub-network centered around protein *STX1A_*1, *SNAP25*_2, *STXBP_* 1 and gene *VAMP2.* They are the major components of the SNARE complex, which medicates the fusion process of synaptic vesicles and play an essential role in the cross talk between neurons and glias [58, 59]. Top pathways enriched by these genes form a functional group related to release neurotransmitter cycle in Fig.3. More specifically, most of them are involved in the GABA synthesis, release, reuptake and degradation (adjusted p=9.532e-11). GABA has been found to have significantly reduced levels in severe cases of AD [60]. Selective inhibition of astrocytic GABA synthesis or release has been suggested as a potential therapeutic strategy for treating memory impairment in AD [61].

### Replication Study on MSBB Data Set

Replication analysis was done using the genotype, RNA-Seq gene expression and protein expression from the MSBB cohort. Due to different protein quantification methods, peptides in MSBB can’t be directly matched to those in ROS/MAP. Therefore, we included all the peptides that belong to the 126 unique proteins in ROS/MAP. Across all the SNPs, genes and proteins used in discovery analysis, we found 107 peptides, 648 genes, and 695 SNPs in 121 MSBB participants. The optimal hyper-parameters learned on training set obtained 82% accuracy on testing data set, close to the test accuracy of MoFNet on ROS/MAP data. We followed the same procedure as discovery analysis to select the optimal cut-off threshold (0.01) and pruned the prior trans-omic network. Finally, 116 out of 313 features identified using ROS/MAP data were replicated, i.e., 56.9% for proteins, 29.6% for genes and 23.6% for SNPs respectively. For the largest connected component (component 1) in Fig. 2(a), the overall replication rate was 48.6%. Specifically, the replication rate is 75% and 64.7% for proteins and genes respectively. For the third connected component, the overall replicated rate was at 50%, with protein replicated at 43% and genes at 50%. For second network component, only 50% SNPs were replicated but not the others. Overall, proteins are more likely to be replicated, which is as expected since proteins integrated information from functionally connected SNPs and genes.

## Discussion

Genes/proteins in the 3rd network component form the core SNARE (soluble N-ethylmaleimide-sensitive factor attachment protein receptor) complex priming the synaptic vesicle fusion for neurotransmitter release, including synaptobrevin (also known as vesicle-associated membrane protein 2 *(VAMP2*)) on the synaptic vesicle, syntaxin 1 *(STX1A/STX1B*) on the plasma membrane, and SNAP25. These three functions together with *STXBP1* to bring the membranes of synaptic vesicle and plasma membrane into apposition and to enable neurotransmission [58]. The SNARE complex is necessary not only for neuron–neuron communication but also neuron–glia and glia–glia communication [59].

Interestingly, for the largest component 1, enrichment results of genes/proteins using EnrichR are also highly relevant to synaptic functions, in addition to signaling pathways and immune system [62]. More specifically, 23 out of 34 genes in component 1 are involved in immune system (adjusted *p* = 2.289*e* – 14), 25 in signaling transduction (adjusted *p* = 1.034*e* – 15), 14 in axon guidance (adjusted *p* = 2.235*e* – 12) and 14 in synaptic function (aggregated from all enriched synaptic GO terms with adjusted *p* ≤ 0.05). Out of 14 synaptic function related genes, 10 of them are involved in both immune system and signal transduction and 7 of them are related to neuron axon guidance. This suggests that synaptic function is an integral part of immune response and signal transduction. Top signaling pathways enriched by component 1 like Notch pathways, NTR (neurotrphin) signaling pathways, and NGF (nerve growth factor) are all activated by their corresponding receptors located in the cell membrane. SNARE complex in component 3 along with GTPase will help merging endocytic vesicles that transport these receptors to the plasma membrane [63], which agrees with previous findings that notch signaling is likely regulated by intracellular vesicle trafficking [63]. These signaling pathways play essential role in normal development of neurons and glias and mediating their cross-talk [64].

Alterations in synaptic function and synapse degeneration are considered among the earliest changes of AD, even before the accumulation of misfolded protein aggregates and neuronal loss [65–67]. Synaptic dysfunction is considered as contributing to both onset and progression of AD and synapse loss in postmortem brains is the strong correlate with cognitive decline [68, 69]. Yet, the connection of synapse dysfunction with amyloid and tau pathology is not fully understood. The synapse is the place where amyloid-beta peptides are generated and is the target of the toxic amyloid-beta oligomers [70]. Oligomeric Amyloid-beta is found present in both pre- and post-synaptic puncta and its co-localization with Apolipoprotein E4 protein is associated with significant increase of Amyloid-beta at synapses [71]. Data from animal models found that Tau binds to synapse vesicle and interferes with presynaptic functions, like synaptic vesicle mobility and neurotransmitter release rate [72]. A recent study reported the interaction of tau in neuron with *STX1A*, a member of the SNARE complex, suggesting the localization of tau at sites of presynaptic vesicle fusion [73].

Taken together, both amyloid and tau are likely to localize at the presynaptic vesicle and have domain-specific interactions with synaptic vesicle-associated proteins, interfering with synaptic vesicle function in the early stage of AD. Also, considering that all these genes are preferably expressed in brain according to GTEx [74] and are mostly markers of astrocyte, microglia and neurons (Fig. 3), this interference of synaptic function could possibly cause disruption in neuron-neuron or neuron-glia cross talk and further lead to neuronal and synapse loss in AD. Further investigation of these identified genes/proteins could possibly help decipher the mechanisms underlying synaptic dysfunction in AD, and ultimately inform therapeutic strategies to modify AD progression.

## Conclusion

We proposed a new deep graph fusion network to leverage the information flow from DNA to proteins such that gene expression and protein expression data can be enhanced with improved prediction power. Known functional connections between SNPs, genes and proteins were embedded into the neural network as prior knowledge. Edge weight and node importance score learned from MoFNet further help prune down the prior network where only sub-networks predictive of the disease outcome will be retained. MoFNet showed superior performance over other integrative -omics models in three ways: 1) it jointly models genotype, gene expression, protein expression and their prior functional relationships, 2) it yields sub-networks predictive of outcomes, instead of individual markers with less interpretability, and 3) it enhances gene expression and protein expression data by leveraging the information flow from DNA to protein. Trans-omic paths from MoFNet findings suggested that AD may be partly the result of genetic variations due to their potential cascading effects on the downstream transcriptome and proteome levels. While none of the prior functional connections was extracted in a tissue specific manner, eQTL analysis showed that MoFNet can accurately pick out those tissue specific relationships between SNPs and genes.

It is worth mentioning that the integrative analysis in this paper is still limited in several ways. First, this is a targeted analysis focusing on only a small set of functionally connected multi-omic features, the number of which was further limited because of the bottleneck in protein expression data. Second, due to the incompleteness of multi-omic data, only a portion of participants were included. Therefore, the classification performance can not reflect the true predictive power of these three types of multi-omic data, which is expected to be much higher if with more -omics features and samples. This study is also limited in that amnestic mild cognitive impairment group (MCI, a transition stage between CN and AD) is excluded from the analysis. How to include this group and investigate the stage-specific multi-omic sub-networks warrant further effort. In addition, like many current multi-omic models for joint analysis, MoFNet requires concatenation of multi-omic features, leading to exclusion of large chunk of samples. An improved version of MoFNet capable of handling the incomplete multi-omic data will also be of great value to the field.

## Author contributions statement

L.X.: Conceptualization, Methodology, Visualization, Formal analysis, Validation, Writing - original draft, Writing review & editing. Y.R.: Conceptualization, Methodology, Formal analysis, Validation, Writing - original draft. P.V.: Conceptualization, Visualization. B.H.: Investigation, Visualization, Formal analysis. S.R.: Data curation, Writing – review & editing. K.N.: Data curation, Writing – review & editing. P.S.: Methodology, Supervision. A.S.: Data curation, Resource, Writing – review & editing. J.Y.: Conceptualization, Methodology, Visualization, Writing - original draft, Writing review & editing, Supervision, Funding acquisition.

## Acknowledgments

The results published here are in whole or in part based on data obtained from the AMP-AD Knowledge Portal. Study data were provided by the Rush Alzheimer’s Disease Center, Rush University Medical Center, Chicago. Data collection was supported through funding by NIA (grants P30AG10161, R01AG15819, R01AG17917, R01AG30146, R01AG36836, U01AG32984, U01AG46152), the Illinois Department of Public Health and the Translational Genomics Research Institute.

## Key points

- An interpretable multi-omic graph neural network model was developed to identify a group of functionally connected multi-omic features associated with Alzheimer’s disease.
- Multi-omic modules identified by proposed model are related to immune system, protein degradation and neurotransmitter release respectively.
- The identified gene and proteins are highly related to synaptic vesicle function.

## Acknowledgments

This research was supported by NIH grants R21 AG066135, R01 EB022574, R01 AG019771, P30 AG010133, NSF CRII 1755836 and NSF CAREER 1942394. The results published here are in whole or in part based on data obtained from the AMP-AD Knowledge Portal. Study data were provided by the Rush Alzheimer’s Disease Center, Rush University Medical Center, Chicago. Data collection was supported through funding by NIA grants P30AG10161, R01AG15819, R01AG17917, R01AG30146, R01AG36836, U01AG32984, U01AG46152, the Illinois Department of Public Health, and the Translational Genomics Research Institute.

## Data Availability

The source code is available through GitHub. Multi-omic data used in this analysis is from the ROS/MAP project and is available after application through the AMP-AD knowledge portal.

## Footnotes

https://adknowledgeportal.synapse.org

https://github.com/JW-Yan/MoFNet

