## Supplementary for "Deep trans-omic network fusion reveals altered synaptic network in Alzheimer’s Disease"

**S.1 GWAS Genotype Data Pre-processing** Imputed GWAS genotype data from the ROS/MAP cohort was downloaded from the AMP-AD knowledge portal. Briefly, raw genotype data underwent the quality control (QC) analysis, as described in ^[1]^. Only individuals with European ancestry were genotyped to minimize population heterogeneity. Sample-level quality control assessment included exclusion of samples with genotype success rate < 95%, discordance between inferred and reported gender, and excess inter/intra heterozygosity. SNP-level quality control assessment included exclusion of SNPs with Hardy-Weinberg equilibrium (*p* < 0.001), MAF < 0.01, genotype call rate < 0.95, mishap test < 1e-8. Population outliers were identified and removed using EIGENSTRAT with default parameters. Genotypes were imputed using BEAGLE software. With the 1000 Genomes Project as the reference panel ^[2]^. AMP-AD portal only provides genetic variants from whole genome sequencing data for MSBB cohort. From that, we extracted the genotype of the same set of SNPs used in discovery analysis. Out of 354 sequenced human DNA samples, 349 of them were included for the subsequent variant calling after removing samples with sequences of low quality and duplicated sequences. All samples have undergone rigorous quality assessment using a comprehensive set of quality measures upon completion of each step of sample processing: 1) sample receipt, 2) library preparation, 3) sequencing, 4) data analysis. Detailed processing steps of MSBB WGS data can be found in ^[3]^.

**S.2 RNA-Seq Data Pre-processing** RNA-Seq gene expression data in ROS/MAP was collected from the dorsal lateral prefrontal cortex (DLPFC) tissue of postmortem brains. It was reprocessed in parallel with several other RNA-Seq data sets from the Accelerating Medicines Partnership for Alzheimer’s Disease (AMP-AD) ^[4]^, including those from the MSBB cohort. For MSBB cohort, processed RNA-Seq data from Brodmann 10 region was downloaded for replication analysis since it is the nearest region to DLPFC, where ROS/MAP RNA-Seq was collected. Brain samples were prepared and sequenced on the Illumina HiSeq 2500 System with 100 nucleotide single end reads, according to the standard manufacturer’s protocol. The raw sequence reads from both cohorts were aligned to the GENCODE24 (GRCh38) reference genome using spliced transcripts alignment to a reference (STAR) ^[5]^. To evaluate the quality of individual samples and identify potentially important covariates for expression modeling, two sets of metrics were calculated using the CollectAlignmentSummaryMetrics and CollectRnaSeqMetrics functions in Picard ^[6]^. Transcript abundances were estimated for each sample using Sailfish ^[7]^. Downloaded RNA-Seq data sets for ROS/MAP and MSBB cohorts were already corrected for known confounding factors, including PMI, RACE, Batch, SEX, RIN and Exonic rate ^[3]^.

The RNA-Seq data from MSBB was collected from Brodmann Areas 10, 22, 36 and 44. The specific brain regions underwent dissected, pulverized stages and stored at cooled environment. RNA samples were isolated at two RNA preparation cores in Mt. Sinai, Human Immune Monitoring Center (HIMC) and quantitative Polymerase Chain Reaction (qPCR). At both cores, the total RNA were isolated from brain tissues using RNeasy Lipid Tissue Mini Kit from Qiagen (cat\#74804) according to the manufacturer’s protocol (The RNeasy Lipid Tissue Mini Kit Handbook, Qiagen 104945, 02/2009) with slight modifications ^[3]^.

**S.3 Proteomic Data Pre-processing** For all ROS/MAP participants, targeted selective reaction monitoring (SRM) proteomics data from dorsal lateral prefrontal cortex tissue (DLPFC) was downloaded from the AMP-AD knowledge portal (10.7303/syn10468856). In total, there are 186 peptides measured corresponding to 126 unique genes. The samples were prepared for LC-SRM analysis using the standard protocol ^[8]^. All the data was manually checked to ensure that the peak assignments and boundaries were valid. The abundance of endogenous peptides was quantified as a ratio to spiked-in synthetic peptides containing stable heavy isotopes. The "light/heavy" ratios were log2 transformed and shifted such that the mean log2-ratio was zero.

Proteomic data from prefrontal cortex brain tissues from Mount Sinai School of Medicine Brain Bank (MSBB) was downloaded from the same portal. It was measured with a different mass spectrometry-based protein quantification approach - Label Free Quantification (LFQ) method. The resulting mass spectrometry data was processed using a common pipeline as detailed in ^[9]^. The processed protein expression data was directly downloaded from the AMP-AD knowledge portal. We included proteins with fewer than 30% missing values in the subsequent analyses ^[3]^. For all the peptides with missing data, imputation was conducted following the data-processing pipeline of probabilistic minimum imputation method ^[10]^.

Finally, we removed the effects of age, gender, post-mortem interval (PMI) and experimental batch using a linear regression model, even though these covariates did not have a strong effect on the data ^9^. Detailed steps regarding brain tissue homogenization, protein digestion, mass spectrometry analysis and quantification for both cohorts can be found in ^[3]^.

### S.4 Parameter Tuning We applied 5-fold cross validation with grid search to tune the parameters given the small sample size. The AD/CN ratio was maintained the same across all training, validation and test partitions. To ensure the fair comparison of competing methods, all methods including the proposed MoFNet went through exactly the same parameter tuning process with 5-fold cross validation. All the training, validating and testing partitions used across methods were kept exactly the same across methods. The search space of L1 regularization rate, learning rate, dropout rate and weight decay rate was set from 0.00001 to 0.01, 0.0001 to 0.01, 0.1 to 0.9, and 0.00001 to 0.001, respectively. The dimension of last 3 fully connected layers were picked from a series of 160 to 8 decremented by 8, and a series of 150 to 10 decremented by 10. Finally, the best performance of MoFNet was obtained with L1 regularization rate as 0.001 applied on the first transparent layer only. The best learning rate as 0.001, dropout rate as 0.5, and weight decay as 0.0008 were applied globally to each layer. The dimension of last 3 fully connected layers are (186, 100), (100,32) and (32,1) respectively. Additionally, we used early termination and dropout layers to avoid overfitting. For the replicated study using MSBB data, since the peptides are collected using different quantification methods, data from two cohorts cannot be directly matched. Therefore, we repeated the parameter tuning process. Given the imbalance of AD and CN groups, random over-sampling on the minority class was applied ^[11]^. Given the small sample size, 3-fold cross validation was applied instead to tune the parameters. We followed the same grid search strategy used in the discovery study. Finally, the best performance of MoFNet was obtained on the replication data, with L1 regularization rate as 0.001 on the first transparent layer, global hyper-parameters learning rate as 0.001, dropout rate as 0.5, and weight decay as 0.0001.

### S.5 Supplementary Figures

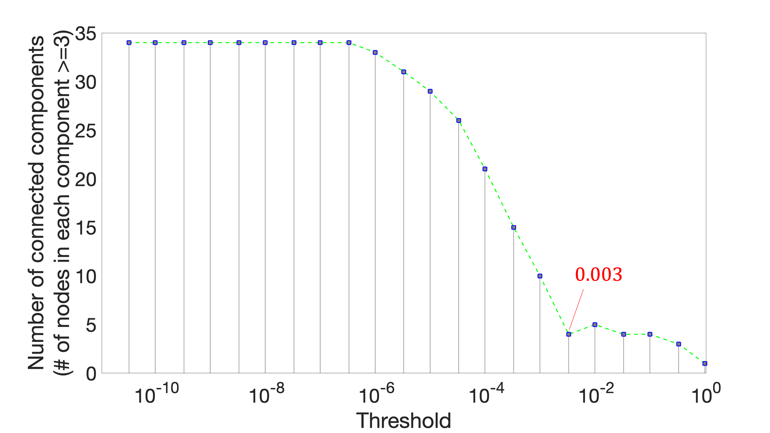

**Figure S1** Selection of cut-off threshold for ROS/MAP data.

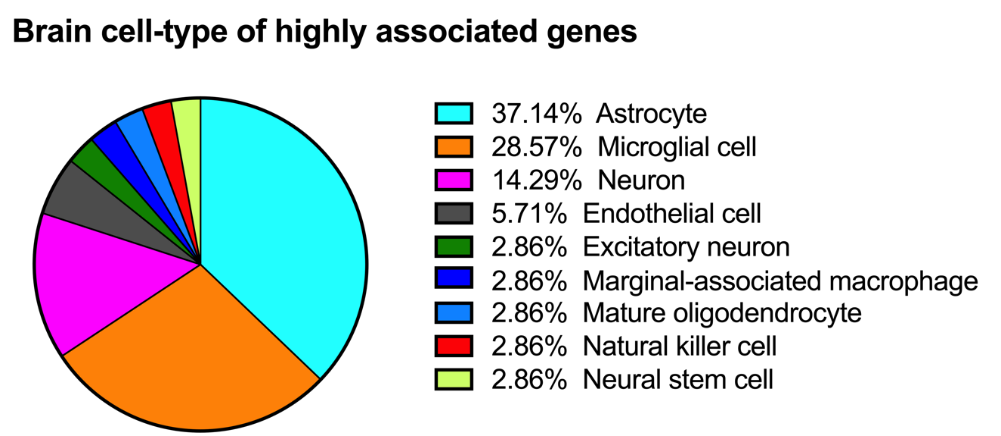

**Figure S2**: Distribution of cell types enriched by the genes and proteins identified in MoFNet.
